# Thinking too positive? Revisiting current methods of population-genetic selection inference

**DOI:** 10.1101/009654

**Authors:** Claudia Bank, Gregory B. Ewing, Anna Ferrer-Admettla, Matthieu Foll, Jeffrey D. Jensen

**Affiliations:** School of Life Sciences, Ecole Polytechnique Fédérale de Lausanne (EPFL), 1015 Lausanne, Switzerland; Swiss Institute of Bioinformatics (SIB), 1015 Lausanne, Switzerland; Department of Biology and Biochemistry, University of Fribourg, 1700 Fribourg, Switzerland

**Keywords:** natural selection, background selection, population-genetic inference, evolution, computational biology

## Abstract

In the age of next-generation sequencing, the availability of increasing amounts and quality of data at decreasing cost ought to allow for a better understanding of how natural selection is shaping the genome than ever before. Yet, alternative forces such as demography and background selection obscure the footprints of positive selection that we would like to identify. Here, we illustrate recent developments in this area, and outline a roadmap for improved selection inference. We argue (1) that the development and obligatory use of advanced simulation tools is necessary for improved identification of selected loci, (2) that genomic information from multiple-time points will enhance the power of inference, and (3) that results from experimental evolution should be utilized to better inform population-genomic studies.

## Identification of beneficial mutations in the genome: an ongoing quest

The identification of genetic variants that confer an advantage to an organism, and that have spread by forces other than chance, remains as an important question in evolutionary biology. Success in this regard will have broad implications not only for informing our view of the process of evolution itself, but also for evolutionary applications ranging from clinical to ecological. Despite the tremendous quantity of polymorphism data now at our fingertips, which in principle ought to allow for a better characterization of such adaptive genetic variants, it remains a challenge to unambiguously identify alleles under selection. This is primarily owing to the difficulty in disentangling the effects of positive selection from those of other factors that shape the composition of genomes, including both demography as well as other selective processes.

Approaches to identify positively selected variants from genomic data can be broadly divided into two categories: those that make use of within-population polymorphism data, and those that make use of between-population/species data. While each approach has its respective merits, we here focus on recent developments in population-genetic inference from polymorphism data in both natural and experimental settings (see [1,2] for more general reviews, and [3,4] for recent and specific literature on divergence-based selection inference). For population-genetic inference from single-time point polymorphism data (as is most commonly the case) this includes not only sophisticated statistical methods, but also simulation programs that enable us to model expected genomic signatures under a wide variety of possible scenarios. Alternatively, data from multiple-time points – such as those recently afforded by ancient genomic data as well as many clinical and experimental datasets - can be used to greatly improve inference by catching a selective sweep “in the act”. Finally, recent results from the experimental evolution literature have begun to better illuminate the expected distribution of fitness effects (DFE), the associated costs of adaptation, and the extent of epistasis. In this opinion piece, we present an overview of recent developments for selection inference in the above-mentioned areas, and offer a roadmap for future method development.

## Selection inference from a single-time point

One of the earliest efforts to quantify selection in a natural population was based on multiple-time point phenotypic data [5,6]. With the onset of genomics however, new sequencing methods were both tedious and expensive, such that the vast majority of data collected were, and still are, of the single-time point variety. For this reason, one generally observes only the footprints of the selection process, making it more difficult to distinguish regions shaped by neutral processes from those shaped by selective processes. Over the past decades, population genetic theory has predicted the effects of different selection models on molecular variation. These predictions have given rise to test statistics designed to detect selection using polymorphism data, based on patterns of population differentiation (e.g., [7-10]), the shape of the site-frequency spectrum (SFS) (e.g., [11-13]), and haplotype / linkage disequilibrium (LD) structure (e.g., [14-17]).

Despite efforts to create statistics robust to demography, all currently available methods to detect selection are prone to misinference under non-equilibrium models [18,19]. Therefore, in parallel to the production of statistics for inferring selection, a separate class of methods has been developed to estimate the demographic history of populations utilizing the same patterns of variation[20,21]. This lends itself to a two-step approach when analyzing population-genetic data: first, demography is inferred using a putatively neutral class of sites, and that model is then used to test for selection among a putatively selected class of sites [13,16]. However, the assumptions enabling this inference are highly problematic because they rely on the correct identification of a class of sites that is both neutral and untouched by linked selection. Background selection (BGS), for example, may indeed influence a large fraction of the genome in many species [22]. Thus, the misidentification of such a class of sites in the genome may not only result in the misinference of the underlying demographic history, but also in misinference of selection owing to the incorrect estimation of the demographic null model – leading to both false positives and false negatives (Figure 1).

**Figure 1:**
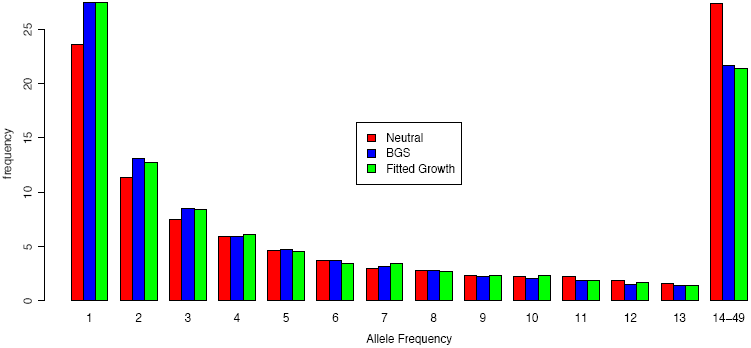
An example of the ability of background selection to mimic neutral non-equilibrium (here, exponential growth) SFS-based patterns. Assuming an absence of BGS effects when they are present (as is common in demographic inference), may result in substantial misinference of the underlying demographic model – here inferring population growth when the population is in fact at equilibrium. Simulations were performed using SFSCode [40], with 50 samples from a single-time point, conditioned on 100 single-locus polymorphisms per locus. The BGS coefficient is ***α****=2Ns=-4* and the probability of a deleterious mutation is 0.1, with recombination rate ***ϱ****=2Nr=50* between chromosome ends. The SFS shows the frequency of occurrence (y-axis) of a number of derived alleles (x-axis) in the simulated sample. Site classes 14-49 were binned for illustrative purposes.

In order to circumvent this problem, it is therefore necessary to develop methods that can jointly infer the demographic and selective history of the population simultaneously, recognizing that both processes are likely to shape the majority of the genome in concert [23]. The development of simulation software that can model both positive and negative selection in non-equilibrium populations (see below) is a step towards this goal, as it allows for the generation of expected patterns of variation under such scenarios. Recently, the introduction of likelihood-free inference frameworks such as Approximate Bayesian Computation (ABC) [24] has made it computationally feasible to combine demographic and selective inference. Recent advances in this field allow for management of the large number of summary statistics needed to achieve this goal [25,26], and the pieces appear largely in place for significant progress to be made on this front.

## Selection inference from multiple-time points

As noted above, although temporal data was considered in the infancy of population genetics, it was not until the late 1980’s that multi-time point genetic datasets became available, spurring the development of related statistics. These approaches focused first on the estimation of changes in population size [27-30]; later, new methods were developed specifically for time-serial data to co-estimate parameters such as the intensity of selection and the effective population size [31].

Multi-time point methods have a major advantage over single-time point based approaches in that knowledge of the trajectory of an allele provides valuable information about the underlying selection coefficient. Owing to advances in sequencing technologies, there is now an increased availability of multi-time point data of not only an experimental but also a non-experimental nature (ranging from ancient samples to those from longitudinal medical studies or field work). This temporal dimension spurred the development of multiple-time point based methods over the last few years – all seeking to estimate some combination of selection coefficient and a) effective population size [32,33], b) migration [34] c) the age of the selected mutation [31], or d) the recombination rate [35]. These methods differ both in the underlying models and in their respective performance, and hence in the conditions to which they are best suited (cf. Table 1, Supplementary Table 1). In Malaspinas *et al.* [31] and Mathieson and McVean [34], the trajectory of the selected allele is modeled as a hidden Markovian process, and the observations are represented as binomial observations from the population. Subsequently, Malaspinas *et al.* [31] use an approximate transition density to compute the likelihood, whereas Mathieson and McVean [34] use an expectation-maximization algorithm to maximize the likelihood, which speeds up the estimation procedure. By contrast, Foll *et al.* [32,33] do not use a Hidden Markov Model (HMM) but an ABC approach to obtain the posterior distribution of the selection coefficient after simulating allele frequency trajectories using the Wright-Fisher model under a range of selection coefficients. This approach results in lower accuracy for very small selection coefficients, but also represents the fastest and least biased method to date.

**Table 1.**
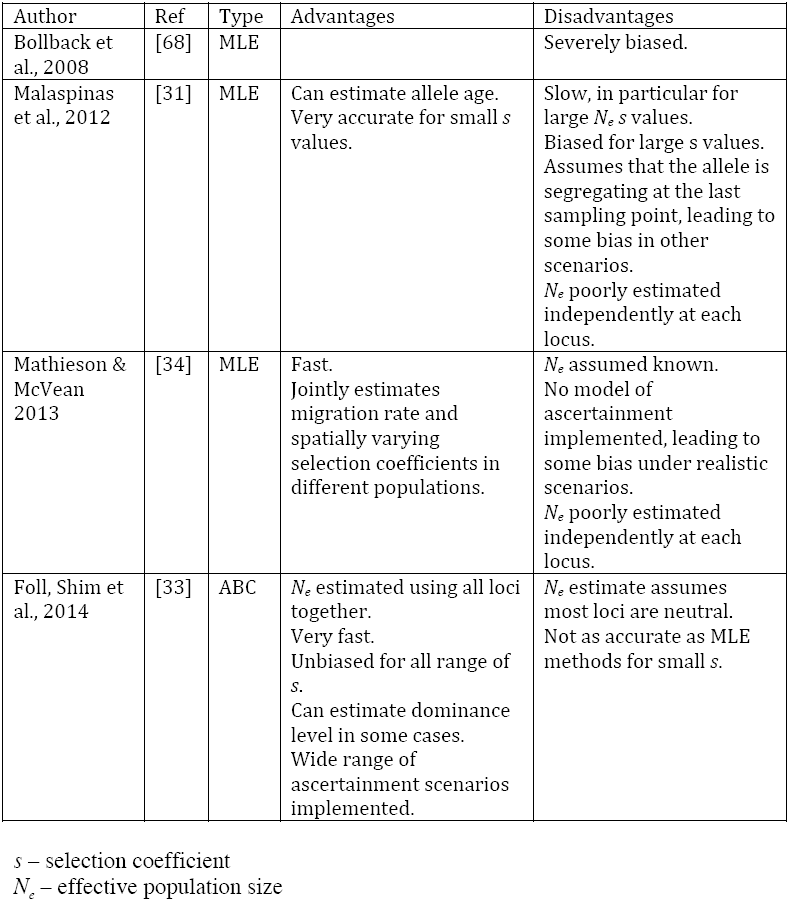
An overview of *currently available multiple-time point selection estimators and their strengths and weaknesses.*

| Author | Ref | Type | Advantages | Disadvantages |
| --- | --- | --- | --- | --- |
| Bollback et al., 2008 | [68] | MLE |  | Severely biased. |
| Malaspinas et al., 2012 | [31] | MLE | Can estimate allele age.<br>Very accurate for small $s$ values. | Slow, in particular for large $N_e s$ values.<br>Biased for large $s$ values.<br>Assumes that the allele is segregating at the last sampling point, leading to some bias in other scenarios.<br>$N_e$ poorly estimated independently at each locus. |
| Mathieson & McVean 2013 | [34] | MLE | Fast.<br>Jointly estimates migration rate and spatially varying selection coefficients in different populations. | $N_e$ assumed known.<br>No model of ascertainment implemented, leading to some bias under realistic scenarios.<br>$N_e$ poorly estimated independently at each locus. |
| Foll, Shim et al., 2014 | [33] | ABC | $N_e$ estimated using all loci together.<br>Very fast.<br>Unbiased for all range of $s$.<br>Can estimate dominance level in some cases.<br>Wide range of ascertainment scenarios implemented. | $N_e$ estimate assumes most loci are neutral.<br>Not as accurate as MLE methods for small $s$. |

$s$ – selection coefficient
$N_e$ – effective population size

Although current multi-time point methods are better at inferring the parameters of positive selection than single-time point methods, a number of challenges remain. Although maximum-likelihood estimators (MLE) may have higher accuracy for small selection coefficients than an ABC-based approach, currently available implementations are extremely computationally intensive and thus cannot be readily used as scans of selection (i.e., a candidate site must be known *a priori* – a generally unlikely scenario), and the number of time points necessary to achieve sufficient power still remains largely unexplored. Also, owing to the recent development of these methods, a relatively small number of selection models have been considered – primarily those of consistent and directional positive or negative selection acting on the candidate site. As such, they suffer from the same limitation described above for single-time point inference in that they are unable to jointly estimate selection and the demographic history of the population. Secondly, such methods have also thus far been unable to effectively tease apart the effects of direct vs. linked selection. On the other hand, given the computational efficiency of recently proposed ABC-based approaches [32,33], it has indeed become possible to effectively utilize such approaches in order to identify candidate sites from whole genome polymorphism data, thereby avoiding the first limitation. Also, because of its efficiency and ability to handle multiple summary statistics, ABC appears as the most promising framework in which to consider more complex selection models under non-equilibrium demographic scenarios.

## Simulating selection

Simulation programs have long been an important tool in population genetics, both for validating theoretical work and for characterizing real data. In the area of selection inference, simulations are used to investigate appropriate critical values of test statistics under non-trivial models and for detecting deviations from the standard neutral model [11,36,37]. Recently, the importance of efficient simulation under complex models has become relevant both for understanding expected patterns of variation and for direct inference using likelihood free ABC [24].

Currently, there are a number of simulation programs available that can simulate a wide range of scenarios and are applicable to different types of models and data (see Table 2). Broadly speaking, simulation programs can be split into two different types. Forward simulators can include a wide range of demographic and selective models [38-43]. However, since every gamete needs to be tracked and every generation simulated, they tend to be computationally costly and thus slow. By contrast, coalescent simulators are very fast as only the sample’s genealogy needs to be considered, and thus only events for the genealogy in question need to be tracked [44-47]. Unfortunately, including arbitrary selection models in coalescent simulations is difficult, because coalescent models rely on the assumption that mutations are independent of the genealogy of the sample – an assumption that is violated in essentially any model including selection. While certain scenarios can be modeled circumventing this problem (such as modeling a selected allele as an additional population connected by migration [48]), the integration of more complex scenarios represents a challenge [48,49].

**Table 2:**
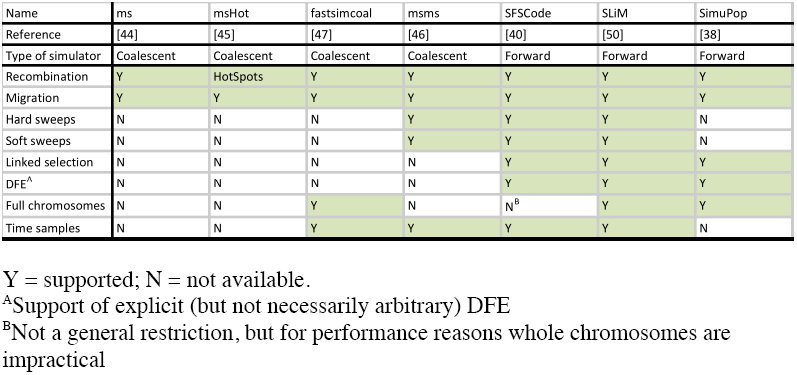
*A selection of commonly used simulation tools and included features.*

| Name | ms | msHot | fastsimcoal | msms | SFSCoDe | SLiM | SimuPop |
| --- | --- | --- | --- | --- | --- | --- | --- |
| Reference | [44] | [45] | [47] | [46] | [40] | [50] | [38] |
| Type of simulator | Coalescent | Coalescent | Coalescent | Coalescent | Forward | Forward | Forward |
| Recombination | Y | HotSpots | Y | Y | Y | Y | Y |
| Migration | Y | Y | Y | Y | Y | Y | Y |
| Hard sweeps | N | N | N | Y | Y | Y | N |
| Soft sweeps | N | N | N | Y | Y | Y | N |
| Linked selection | N | N | N | N | Y | Y | Y |
| DFE<sup>A</sup> | N | N | N | N | Y | Y | Y |
| Full chromosomes | N | N | Y | N | N<sup>B</sup> | Y | Y |
| Time samples | N | N | Y | Y | Y | Y | N |

Y = supported; N = not available.
<sup>A</sup>Support of explicit (but not necessarily arbitrary) DFE
<sup>B</sup>Not a general restriction, but for performance reasons whole chromosomes are impractical

In the development of both forward and backward simulation tools, a major recent focus has been on modeling positive selection under arbitrary non-equilibrium models. As it is difficult to analytically obtain expectations for patterns of variation under these conditions, simulations become important: these models are indeed tractable to simulate, and both forward [50] and coalescent simulation programs [46] have recently been released that provide better performance under models of arbitrary demographic history. Utilizing today’s computational resources, application of more complex models is feasible, allowing for highly parameterized demographic histories [32,51,52], and permitting direct inference of selection using ABC.

More recently, there is growing awareness of the effects of BGS [53,54] in shaping genomic patterns and on the performance of methods designed for detecting positive selection. Intermediate levels of BGS pose a difficult problem in modeling, at least with coalescent simulators, despite the relative ease of simulating BGS in a forward-simulation framework [40]. While strongly deleterious mutations are purged from the population immediately, and very slightly deleterious mutations behave neutrally, intermediately deleterious mutations (cf. Figure 2) accumulate according to non-neutral dynamics that have yet to be studied in more detail [55]. The growing awareness of the importance of accounting for BGS in future studies is comparable to recent progress in the consideration of demography – where previously it was common to assume an equilibrium population history when performing tests of selection, and now demography is commonly considered in such inference. BGS thus appears to be the next frontier to be considered in the development of simulation tools. However, as computing speed and algorithmic developments improve, it should become a priority to develop full inference schemes for co-estimation of demography and BGS in the future [56]. In particular, direct estimation of the DFE in both experimental and natural populations would lead to new insights into the evolutionary process. Variation of estimated parameters over the genome, and when applicable, over time, can lead to a deeper understanding of how the dynamics of demography, recombination, selection and mutation drive evolution in natural populations. Complementary to such inference is the potential for validation of statistical methods under known demographic and selection models via experimental-evolution studies.

**Figure 2:**
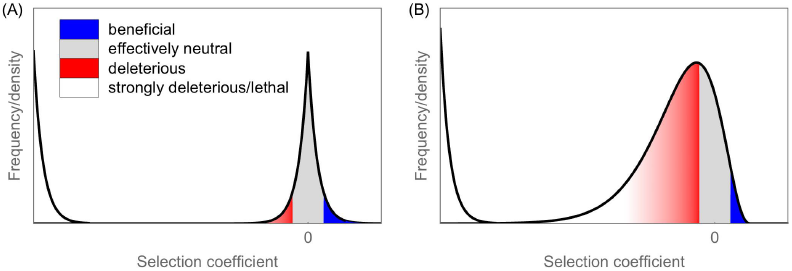
Two hypothetical DFEs that result in different expectations regarding the importance of background selection. (A) A sharp mode around neutrality (here represented as exponential decay in both the negative and positive direction), as frequently assumed in population-genomic studies of the DFE (e.g., [76]), results in little potential for BGS (indicated by the red area under the curve). (B) Experimental-evolution studies of the DFE suggest a wider mode of the DFE around neutrality that is skewed (and sometimes shifted) towards deleterious mutations (here represented by a shifted negative gamma distribution as suggested in [77]). This results in a much higher proportion of slightly deleterious mutations that can contribute to BGS. The intensity of the red shading indicates which area under the curve is most likely to contribute to relevant effects of BGS (see also [66]). The second, strongly deleterious mode of the DFE is depicted as exponentially distributed. This is an arbitrary choice, because the accuracy of currently available methods is not sufficient to detect its actual shape, which should, however, be of little importance due to the severity of the deleterious effects of mutations in that mode.

## Experimental evolution as a new source of information

Through advances in sequencing techniques and bioengineering, experimental-evolution approaches (see Box 1) have become a flourishing area of development in evolutionary biology, catching increasing interest from the field and enabling a large-scale evaluation of mutational effects that was previously unthinkable. Two general types of approaches dominate this area, one relying on the accumulation of naturally arising mutations over many generations, and one based on the simultaneous introduction and competition of hundreds or thousands of engineered mutations. Below, we highlight three implications emerging from recent studies that we believe are particularly relevant for those interested in selection inference.

#### Box 1: Experimental-evolution approaches

In traditional experimental-evolution studies, a lab strain of an organism (typically a virus, bacterium, microbe, or fly) is exposed to a new and potentially challenging environment, and after a number of generations, its fitness (often measured as relative growth rate in competition with the original wild type) is evaluated. The first laboratory evolution experiment of that kind was performed by William Henry Dallinger between 1880 and 1886 – growing microbes in an incubator under increasing temperature, and observing that they evolved to tolerate previously lethal conditions (reviewed in[69]). Today’s approaches fall into two categories that diverge from the original experiment in their increasing use of biotechnology and next-generation sequencing:

1. **Mutation-accumulation experiments**. With the most prominent being Lenski’s long-term selection experiment in E. coli (e.g., [70]) that has been running for more than 50,000 generations, these experiments are very similar to the one explained above, now with the added perspective of whole-genome sequencing that enables identification of mutations accumulated or segregating, and with robots enabling maintenance of hundreds of replicates (reviewed in [71,72]).
2. **Mutagenesis experiments**. Here, hundreds or thousands of individuals are created that carry only one or a few (random or specific) mutations, usually only in a single gene (reviewed in [73]). These are grown for a short period of time and fitness is assessed either by sequencing and estimation of relative growth rate (e.g., [58]), or by assessing other fitness-related phenotypes (e.g., [74,75]).

**Advantages and limitations.** Experimental-evolution studies offer not only new insights into mutational effects, but also a systematic way of testing population-genetic models and theories under controlled conditions – hence bridging theory and nature. They are ideally suited to inform us about expected statistical patterns of the DFE and to study resistance evolution. On the other hand, they are restricted to certain organisms, and it is generally impossible to reproduce the ecology of a natural environment in the lab.

Firstly, approaches using engineered mutations in viruses or microbes allow for a systematic picture of the DFE of all possible point mutations in a chosen region of a gene, hence enabling an assessment of the proportions of beneficial, neutral, deleterious, and even lethal mutations (e.g., [57,58]). The general picture thus far confirms the one proposed by the nearly-neutral theory [59], postulating a bimodal DFE with one mode centered at wild-type-like, and another at strongly deleterious fitness. Notably, however, most studies identify a wide wild-type-like mode strongly skewed towards deleterious selection coefficients (cf. Figure 2). This suggests the heavy influence of effective population size in dictating the neutral class of sites, and further highlights the importance of considering the effects of BGS. Integrating our emerging knowledge of the shape of the DFE into simulation tools will be an important step towards better selection inference.

Secondly, as the scale of experimental-evolution experiments increases, it is more feasible to study not only the effects of single, but also those of double and multiple-step mutations. A pattern of common epistasis emerges (in particular within genes or pathways), in which the combined effects of mutations are very difficult to predict from the measured singular effects of a given mutation. This might result in the “missing heritability” commonly observed in genome-wide association studies [60]. In particular, this suggests that the genetic background must be considered when trying to detect single alleles under selection.

Thirdly, although lab environments cannot reproduce natural conditions (cf., e.g. [61]), there are numerous cases in which laboratory evolution experiments have re-identified known antibiotic or antiviral resistance mutations from natural populations, even if grown under quite unnatural conditions (e.g., [32]), or have provided other important results regarding the evolution of resistance (reviewed in [62]) – indicating that experimental-evolution results are indeed informative for inference about natural populations. In addition, experimental-evolution approaches allow for the study of the effects of the same mutations in different environments. Several such studies have indicated pervasive costs of adaptation, including both mutations of large effect (reviewed in [63,64]) and of small effect [65]. These results suggest that if, upon a change of the environment, adaptation is common from standing genetic variation instead of new mutations, the variation may likely be rare and segregating under mutation-selection balance [66].

## A roadmap for improved selection inference

Despite over one hundred years of population-genetic research, major advances in our understanding of genetics, and huge amounts of polymorphism data across organisms and populations, many of the initial challenges that faced Fisher and Wright still persist today. However, it is fair to say that many of the necessary next steps are clear, and encouragingly, many of the essential pieces appear ready to be assembled to make such advances. We suggest the following as guidelines for moving the field forward:

1. Continued improvements of simulation tools are allowing increasingly relevant models to be considered – but computationally efficient programs capable of simulating a wide range of complex models represent an ongoing challenge. Relatedly, simulations should become an obligatory instrument to assess the power of inference, to test new methods, and to consider alternative models.
2. ABC is a fast and efficient inference method that is likely the way forward for co-estimating demography and BGS with positive selection. However, the identification of summary statistics that are capable of distinguishing between these processes, particularly given that there may not be genomic sites impacted by only a single one of these effects, remains a pressing challenge.
3. Results from experimental evolution complement population genomics by yielding important insights into the DFE and the extent of epistatic interactions. An explicit and realistic shape of the DFE can now be incorporated both into simulation algorithms and in inference methods applied to genomic polymorphism data in natural populations.

This opinion piece thus represents a roadmap for improved selection inference – utilizing novel theory/method development combined with the tremendous amount of data currently being made available via next-generation sequencing. As such, we may for example draw information from experimental-evolution studies, and leverage it against time-sampled population data allowing for entire mutational trajectories to be studied within the context of the underlying distribution of fitness effects. With these theoretical and experimental tools, population genetics is indeed on the verge of important breakthroughs in not only our general understanding of the process of adaptation, but also in the ability of evolutionary analysis to provide important insights to related fields ranging from ecology to medicine.

## Acknowledgements

Financial support for this work was provided by grants from the Swiss National Science Foundation and the European Research Council to J.D.J. The authors are grateful to H. Shim for help with the comparison of multi-time point methods, and to K. Irwin for a careful reading of the manuscript.

**Abbreviations**
ABC: Approximate Bayesian computation
BGS: Background selection
DFE: Distribution of fitness effects
HMM: Hidden Markov model
MLE: Maximum-likelihood estimator
SFS: Site-frequency spectrum

## Glossary

Background selection: reduction of genetic diversity due to selection against deleterious mutations at linked sites.
Coalescent simulator: simulation tool that reconstructs the genealogical history of a sample backwards in time. This greatly reduces computational effort, but only models in which mutations are independent of the sample’s genealogy can be implemented.
Cost of adaptation: the deleterious effect that a beneficial mutation can have in a different environment. Prominent examples are antibiotic resistance mutations, which have often been observed to cause reduced growth rates (as compared with the wild type) in the absence of antibiotics.
Demographic history: the population history of a sample of individuals, which can include population size changes, differing sex ratios, migration rates, splitting and reconnection of the population, as well as variation over time in these parameters.
Distribution of fitness effects (DFE): The statistical distribution of selection coefficients of all possible new mutations, as compared with a reference genotype.
Epistasis: the interaction of mutational effects, resulting in a dependence of the effect of a mutation on the background it appears in.
Forward simulator: simulation tool that models the evolution of populations forward in time. This allows for implementation of complex models, but also usually results in much longer computation times because all individuals/haplotypes must be tracked.
Non-equilibrium model: any model that incorporates violations of the assumptions of the standard neutral model (see below).
Selection coefficient: A measure of the strength of selection on a selected genotype. Usually, the selection coefficient is measured as the relative difference between the reproductive success of the selected and the ancestral genotypes.
Selective sweep: the process of a beneficial mutation (and its closely linked chromosomal vicinity) being driven (“swept”) to high frequency or fixation by natural selection. Selective sweeps result in a genomic signature including a local reduction in genetic variation, and skews in the SFS.
Standard neutral model: under this model [67], the population resides in an equilibrium of allele frequencies determined by the (constant) mutation rate and population size. The model assumptions include random mating, binomial sampling of offspring, and no selection.

